# A targetable N-terminal motif orchestrates α-Synuclein oligomer to fibril conversion

**DOI:** 10.1101/2023.02.10.527650

**Authors:** Jaime Santos, Jorge Cuellar, Irantzu Pallarès, Emily J Byrd, Alons Lends, Fernando Moro, Muhammed Bilal Abdul-Shukkoor, Jordi Pujols, Lorea Velasco-Carneros, Frank Sobott, Daniel E Otzen, Antonio N Calabrese, Arturo Muga, Jan S Pedersen, Antoine Loquet, Jose María Valpuesta, Sheena E Radford, Salvador Ventura

## Abstract

Oligomeric species populated during α-synuclein aggregation are considered key drivers of neurodegeneration in Parkinson’s disease. However, the development of oligomer-targeting therapeutics is constrained by our limited knowledge of their structure and the molecular determinants driving their conversion to fibrils. PSMα3 is a nanomolar peptide binder of α-synuclein oligomers that inhibits aggregation by blocking oligomer to fibril conversion. Here, we investigate the binding of PSMα3 to α-synuclein oligomers to discover the mechanistic basis of this protective activity. We find that PSMα3 selectively targets an α-synuclein N-terminal motif (residues 36-61) that populates a distinct conformation in the monomeric and oligomeric states. This α-synuclein region plays a pivotal role in oligomer to fibril conversion, as its absence renders the central NAC domain insufficient to prompt this structural transition. The hereditary mutation G51D, associated with early-onset Parkinson’s disease, causes a conformational fluctuation in this region, leading to delayed oligomer to fibril conversion and an accumulation of oligomers that are resistant to remodeling by molecular chaperones. Overall, our findings unveil a new targetable region in α-synuclein oligomers, advance our comprehension of oligomer to amyloid fibril conversion and reveal a new facet of α-synuclein pathogenic mutations.

## INTRODUCTION

The aggregation of α-synuclein (αS), a 140-residue intrinsically disordered protein, is a defining hallmark of Parkinson’s disease and related synucleinopathies^1–3^. In these disorders, αS self-assembles into amyloid fibrils that accumulate in the brain of patients, forming insoluble deposits known as Lewy bodies and Lewy neurites. The aggregation landscape of αS is dynamic, involving the formation of transient oligomeric species that precede and co-exist with the final amyloid fibrils^4–9^. αS oligomers are non-fibrillar soluble species that act as key kinetic intermediates in amyloid formation6 and contribute to gain-of-toxic interactions and disruption of cellular processes^10,11^. Therefore, αS oligomers emerge as promising targets for therapeutic and diagnostic interventions^12^, particularly during the early stages of the disease.

Over the past decade, there has been a growing interest in unravelling the structure, formation, and conversion to fibrils of αS oligomers, taking advantage of the ability to kinetically trap these species^8,13–15^. Yet, their highly dynamic nature^14^ poses a technical limit for structural investigations, ultimately hampering the advancement of oligomer-targeting therapies. This emphasizes the need for alternative strategies to investigate the conformational and kinetic properties of αS oligomers.

Phenol-soluble modulin α3 (PSMα3) is a 22-residue amphipathic α-helical peptide that binds αS oligomers with low nanomolar affinity and a 1:1 (αS:PSMα3) stoichiometry^16^. The tight binding of PSMα3 to oligomers contrasts with the lack of any detectable interaction with monomeric αS, underscoring the existence of an oligomer-specific binding site for this peptide (**Figure 1a**). PSMα3 binding abrogates oligomer-associated neurotoxicity and inhibits αS aggregation by blocking oligomer to fibril conversion^16^, thereby interfering with molecular events crucial for pathogenesis. These findings suggest that PSMα3 binding to αS oligomers may be mediated by a therapeutically relevant, oligomer-specific motif.

**Figure 1.**
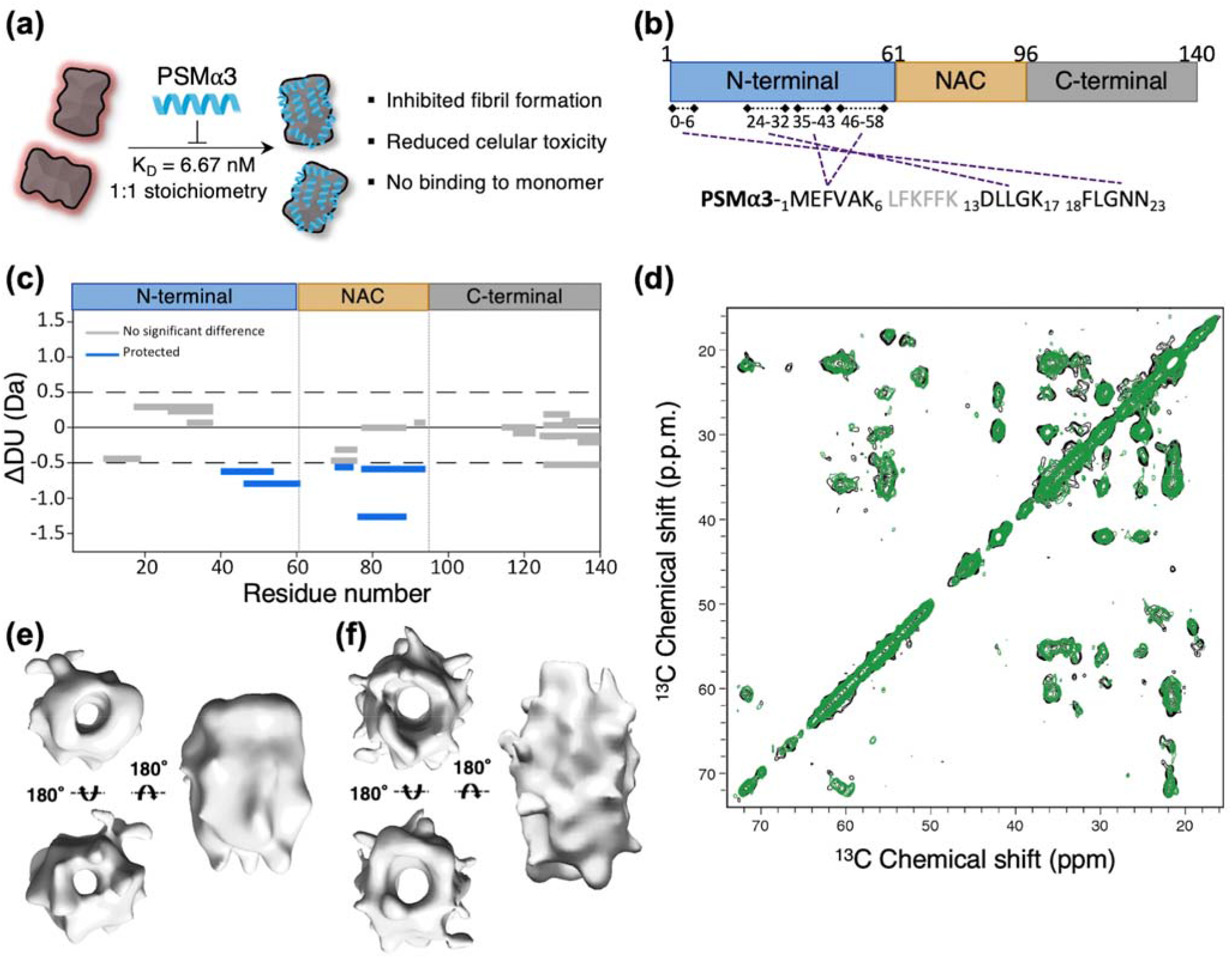
PSMα3 binding to αS oligomers. (a) Schematic representation of PSMα3 binding and activities. (b) Crosslinking map representing PSMα3 contacts with αS oligomers. (c) Wood’s plots showing the difference in deuterium uptake (ΔDU) when comparing αS oligomers in complex with PSMα3 and free αS oligomers by HDX-MS at the 60 second exposure time point. Peptides colored in blue are protected from exchange in presence of PSMα3 (see Methods), suggesting they are less solvent-exposed and/or participate in more inter/intra-protein hydrogen-bonding in the presence of PSMα3. (d) 2D 13C-13C PDSD correlation spectra (mixing time of 50 ms) of oligomers (black) and oligomers + PSMα3 (green). (e) 3D reconstruction of αS oligomers in the absence of PSMα3 (18.5 Å resolution). (f) 3D reconstruction of αS oligomers in complex with PSMα3 (19 Å resolution).

Driven by this idea, here we characterized PSMα3 binding to αS oligomers to identify a new oligomer-specific region implicated in αS pathogenesis. By combining an array of structural, biophysical and biochemical approaches, we found that PSMα3 interacts primarily with a discrete binding site within the N-terminal region of αS, encompassing residues 36-61, which overlaps with two regions (P1 and P2) reported to be ‘master controllers’ of αS aggregation^17–20^. We characterized the role of P1 and P2 in the context of αS oligomers and found that these regions are critical for the oligomer to fibril transition. This structural conversion process is tightly regulated by the sequence of this N-terminal region of αS. Accordingly, we show that a familial mutation within this region associated with early-onset Parkinson’s (G51D) causes a local conformational fluctuation that delays oligomer to fibril conversion, resulting in the accumulation of oligomers that resist disaggregation by molecular chaperones.

Overall, we here identify a disease-relevant αS region fundamental for oligomer to fibril conversion. This sequence defines an oligomer-specific motif that can be targeted by molecular ligands, revealing an uncharted territory for the design of oligomer-directed therapeutic and diagnostic tools.

## RESULTS

### PSMα3 binds to a defined motif in the N-terminal of αS oligomers

To identify the PSMα3 binding site in αS oligomers, we first investigated αS-PSMα3 interactions using crosslinking mass spectrometry (XL-MS) and the zero-length cross-linker, DMTMM (4-(4,6-dimethoxy-1,3,5-triazin-2-yl)-4-methyl-morpholinium chloride). Isolated oligomers were prepared as previously described^13^, incubating 800 μM of monomeric αS for 20 hours at 37 °C quiescently, followed by centrifugation-based fractionation. Oligomer-PSMα3 complexes were generated by incubating oligomers with a 3-fold molar excess of PSMα3 and subsequently removing the unbound peptide by centrifugal filtration. DMTMM crosslinking of the oligomer-PSMα3 complexes revealed four different contact sites between PSMα3 and the N-terminal domain of αS, encompassing residues 1-6, 24-32, 35-43 and 46-58 (**Figure 1b**).

We next sought to confirm the principal segments defining the PSMα3-oligomer interface using hydrogen-deuterium exchange mass spectrometry (HDX-MS). Solvent-exposed residues lacking protein-protein hydrogen bonds incorporate deuterium more rapidly than buried residues or those engaged in inter/intra-molecular hydrogen bonding. Thus, αS amino acids contributing to PSMα3 binding site should exhibit lower deuterium uptake when PSMα3 is bound. We used differential HDX-MS to compare the extent of deuterium incorporation in αS oligomers in the absence and presence of PSMα3. In the presence of PSMα3, two αS peptides covering residues 40 to 61 showed significant protection from deuterium uptake (**Figure 1c, Figure S1**). Together with the XL-MS contacts, this suggests that this N-terminal region constitutes the primary PSMα3 binding site within αS oligomers. Additionally, significant protection from deuterium uptake was identified in three peptides in the NAC domain. Considering the lack of cross-links in this segment, this observation suggests that PSMα3 binding induces a conformational rearrangement of the NAC region which causes a change in solvent accessibility and/or hydrogen bonding in this region.

To address this question, we also investigated the interaction between PSMα3 and αS oligomers using Magic-Angle Spinning (MAS) solid-state nuclear magnetic resonance (ssNMR). MAS-ssNMR has been used previously to define the rigid core of these αS oligomers, comprising residues 70 to 88 within the NAC region^14^. More mobile segments were assigned spanning residues 1-20 and 90-140^14^. Thus, even if the primary PSMα3 binding site (region 35-61, as determined by XL-MS/HDX-MS, **Figure 1b and c**) cannot be assessed by ssNMR, this technique provides a mean to detect conformational shifts within the oligomer’s rigid core upon peptide binding, validating our observations from HDX-MS (**Figure 1c**). We recorded Cross-Polarization (CP) and Insensitive Nuclei Enhanced by Polarization Transfer (INEPT) experiments to measure the ^13^C signals in rigid and mobile segments of αS oligomers, respectively, in the absence or presence of PSMα3. The chemical shifts in the CP-based 2D spectra were identical in both cases (**Figure 1d**). This unique set of resonances was assigned as residues 70 to 89 based on previous studies of αS fibrils (**Table S1** and **Figure S2a**)^21^, and in agreement with those documented for a previously characterized toxic αS oligomer by ssNMR^14^. Given that no chemical shift differences were observed in this region in the presence of PSMα3, this suggests that the protection in this region from deuterium uptake in the presence of PSMα3 detected by HDX-MS is not a result of direct binding or a major structural reorganization of the core. Similarly, only minor chemical shift differences were detected in the INEPT-based 2D spectra (reporting on mobile αS residues 1-20 and 90-140) (**Figure S2b**), consistent with the principal PSMα3 binding site encompassing residues 40 to 61, which are in an intermediate motional regime not accessible to CP and INEPT. Despite the absence of significant chemical shift differences, we noted a clear increase in the CP signal and CP/INEPT ratio in the presence of PSMα3, while sharing the same water hydration dynamics (**Figure S2c-e**). This implies a rigidification or loss of dynamic excursions of the oligomer’s ssNMR-detectable residues upon peptide binding. Considering this, it is conceivable that the protection from deuterium exchange observed in the NAC region of oligomeric αS in the presence of PSMα3 stems from a binding-induced increase in rigidity.

Consistent with the ssNMR data, PSMα3 binding reduced the conformational heterogeneity of αS oligomers, relative to the oligomers alone, as judged by negative stain transmission electron microscopy (nsEM) images (**Figure S3a and b**). This was further sustained by cryo-electron microscopy (cryoEM) 3D density reconstruction of the two sets of particles, revealing a cylindrical architecture with a central hollow core (**Figure 1e and F, Figure S3c-e**), consistent with previous observations of αS oligomers^13^. While the overall architecture of the αS oligomers remained unaltered upon PSMα3 binding, the associated rigidification effect promoted an increase in structural order in the PSMα3-oligomer complexes, noticeable in the 2D classes generated during 3D reconstruction (**Figure S3c-e**). End-on views showed a 6-fold symmetry, also visible in the 3D reconstruction of oligomers, both in absence and presence of PSMα3.

Overall, our data indicate that PSMα3 binds to a specific site within the N-terminal domain of αS, spanning residues 36 to 61. In addition, PSMα3 binding rigidifies, but does not cause a significant structural reconfiguration in the oligomer rigid core.

### The PSMα3 binding site in αS oligomers is partially collapsed and solvent accessible

To gain further insights into the conformation and dynamic properties of the different regions in oligomeric αS, we analyzed the differential deuterium uptake between αS monomers and oligomers using HDX-MS. Peptides in both the N-terminal region and NAC domain become protected upon oligomer formation (**Figure 2a, Figure S1**), whereas no differences in deuterium incorporation in the C-terminal region were observed when comparing αS monomer and oligomers, suggesting that this region remains flexible and disordered in the assembled state. The enhanced protection in the NAC domain upon oligomer formation was expected, considering that this region forms the rigid and structured core of the oligomers according to the ssNMR CP data. Remarkably, the degree of protection was greater for peptides in the N-terminal region compared with those in the NAC domain, suggesting that this N-terminal segment undergoes a significant structural remodeling from the initially disordered state of the monomer. We next applied XL-MS using DMTMM to identify αS to αS contacts within the oligomer. These results confirmed that the enhanced protection from deuterium exchange in the N-terminal domain coincides with the formation of contacts both between different residues within this domain (intra-domain), as well as interdomain interactions between N-terminal and NAC regions (**Figure S4**).

**Figure 2.**
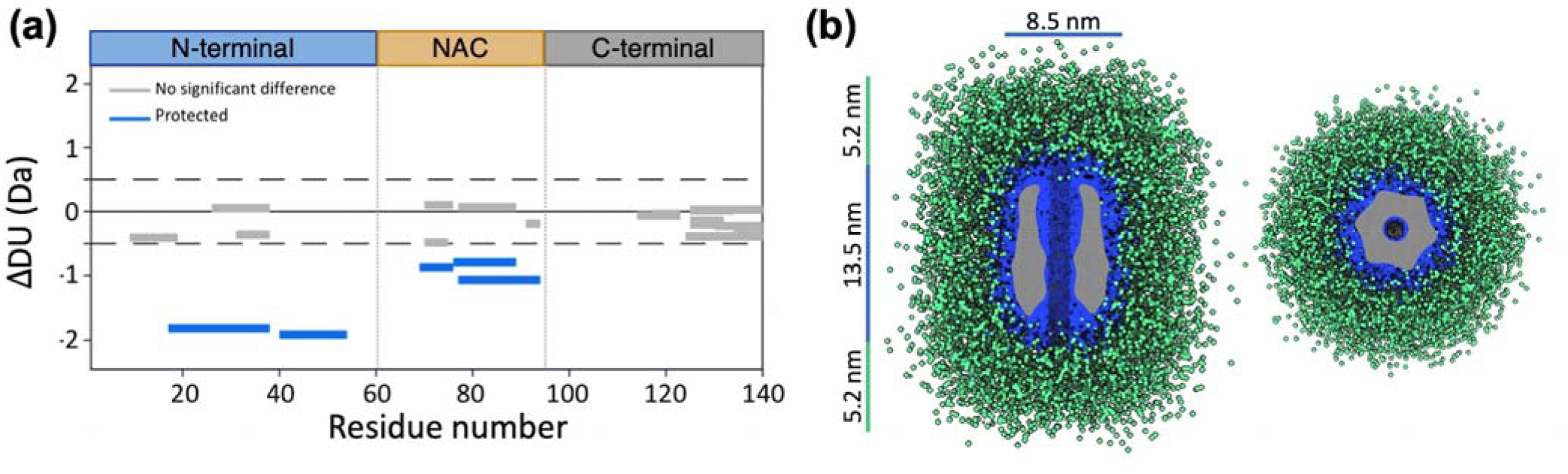
Dynamics of N-terminal region of aS in the oligomer. (a) Wood’s plots showing the difference in deuterium uptake (ΔDU) between αS monomers and oligomers by HDX-MS at the 60 seconds exposure timepoint to deuterium. Peptides colored in blue are significantly protected from exchange in αS oligomers compared with monomeric αS. (b) Two views of the SAXS-based 3D reconstruction of αS oligomers. The compact core (blue) is surrounded by an outer disordered shell (green). The cryoEM density map is shown inside the oligomer core (gray).

Finally, we used small-angle X-ray scattering (SAXS) to probe the conformational properties of the dynamic and disordered αS regions in the oligomer (**Figure 2b; Figure S5a and Table S2**). The oligomer compact core (blue) was modelled as a super-ellipsoid with a central cylindrical hole and showed dimensions (**Table S2**) that align with the cryoEM visible map (gray). This central region is surrounded by an outer shell of disordered tails (green), resulting in an oligomer radius of gyration of 76.1 ± 0.3 Å and dimensions consistent with previous reports (**Figure 2b**)^8,22^. Notably, our experimental SAXS data fitted better to a model with a single random coil chain per monomer rather than two (**Figure S5b**), as would be anticipated if both the N- and C-terminal domains remained fully disordered. In this way, the disordered fuzzy coat was modeled to encompass 48% of αS residues (**Table S2**), whereas the complete N- and C-terminal αS domains account for 76% of the αS sequence. In contrast, the principal contributors to the oligomer INEPT spectra^14^ (residues 1-20 and 90-140) account for 50% of the αS sequence.

Together, these results indicate that the primary constituents of the outer disordered corona of the oligomers are the C-termini, with a contribution of the 20 N-terminal residues of αS. Other residues in the N-terminal domain, containing the PSMα3 binding site, presumably form a partially structured or collapsed conformation that is sufficiently dynamic to be undetectable in the ssNMR CP spectra. The high affinity of PSMα3 for this region and the 1:1 (αS:PSMα3) stoichiometry underscore specificity and accessibility of these N-terminal segments for interactions within the oligomer assembly.

### The N-terminal P1 and P2 regions of αS control oligomer-to-fibril conversion

The PSMα3 binding site in the αS oligomers encompasses two regions in the N-terminal of αS known to be pivotal for amyloid formation, P1 (36-42) and P2 (45-57)^17,18^. These sequences act as ‘master controllers’ of αS amyloid formation *in vitro* and in *Caenorhabditis elegans*, with deletions or point mutations in P1 and P2 suppressing or delaying amyloid formation ^17,18^. Since binding of PSMα3 blocks oligomer to fibril transition^16^, we investigated the specific contributions of P1 and P2 to this conformational conversion.

We assayed the impact of P1 and P2 deletions (ΔP1 and ΔP2) on αS amyloid formation using thioflavin-T (Th-T) fluorescence as a reporter. In agreement with previous results, deletion of P1 (ΔP1) inhibited αS amyloid formation, whereas the deletion of P2 (ΔP2) retarded aggregation under the conditions used^17,18^ (**Figure 3a**). nsEM of the end-point of the aggregation reaction confirmed that wild-type (WT) αS formed mature amyloid fibrils. In contrast, none or few fibrils were present in the ΔP1 and ΔP2 variants at the end-point of the reactions (**Figure S6**). We then applied a centrifugation-based protocol to study the presence of oligomeric species in those samples (see Methods section). As expected, few low molecular weight species (non sedimentable particles with a molecular weight >100 kDa) were visible at the endpoint of the WT αS amyloid assembly reaction (**Figure 3b**). In contrast, the primary components in the low-molecular weight fraction for ΔP1 and ΔP2 inhibited reactions were oligomers identical in shape and size to WT oligomers (**Figure 3b**). Similar results were obtained when both P1 and P2 were deleted in tandem (ΔΔ) (**Figure 3a and b, Figure S7**). These findings indicate that while P1 and P2 are not necessary for αS oligomerization, they actively contribute to the conversion of oligomers into fibrils.

**Figure 3.**
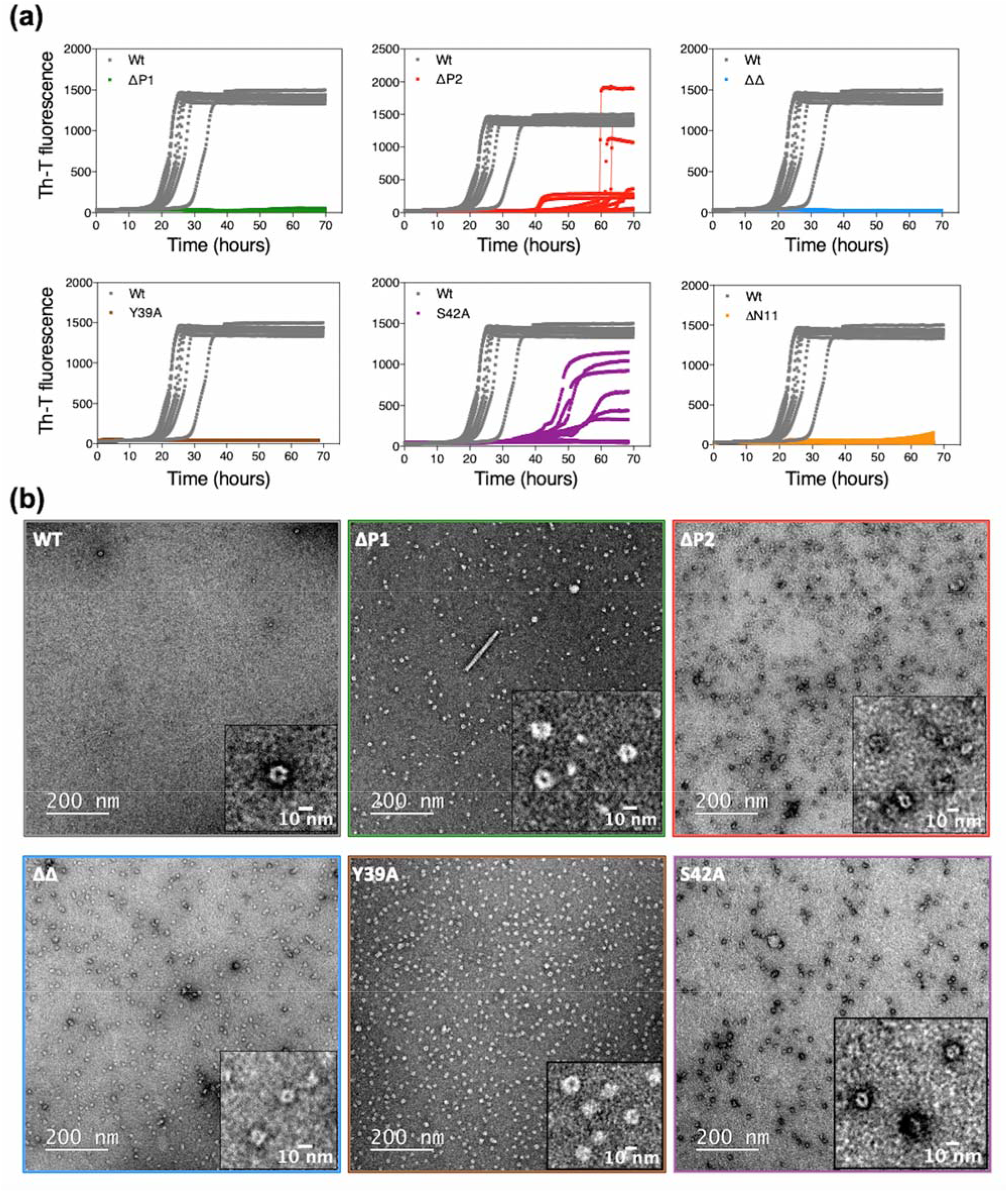
Contribution of PSMα3 binding site to oligomer to fibril conversion. (a) Kinetics of amyloid formation of the WT, ΔP1, ΔP2, ΔΔ, Y39A, S42A and ΔN11 variants monitored using Th-T fluorescence. (b) Representative nsEM images of the oligomeric fraction of the WT (top left), ΔP1 (top middle), ΔP2 (top right), ΔΔ (bottom left), Y39A (bottom middle), and S42A (bottom right) isolated at the endpoint (WT, ΔP1, ΔP2, ΔΔ, Y39A) or after 28 hours of assembly (S42A).

The modulation of amyloid formation by the P1 region has been shown to be dependent on specific residues^18^. Hence, we characterized two αS variants, Y39A and S42A, which were shown previously to mimic the ΔP1 variant, inhibiting amyloid fibril formation^18^. Consistent with prior results, the Y39A and S42A amino acid substitutions within the P1 region inhibited αS amyloid formation to different extents (**Figure 3a, Figure S7**). In both cases, oligomers akin to those observed above for WT αS were the predominant species in the oligomeric fraction at the time points of maximal inhibition (end point for ΔP1 and 28 hours for ΔP2), indicating an impaired transition of oligomers to fibrils (**Figure 3b**). Notably, the WT αS and S42A proteins differ by just a single hydroxymethyl group, evidencing the precise control that subtle sequence changes can exert on the conversion of oligomers to fibrils.

To determine whether oligomer accumulation is specific to sequence changes or deletions in the P1 and P2 regions, we characterized an N-terminal truncated variant (ΔN11) which has been shown to result in delayed amyloid assembly due to decreased secondary nucleation^7,23^. As expected, deletion of the N-terminal eleven residues inhibited amyloid formation, but oligomers were barely detectable in the low-molecular weight fraction (**Figure 3a, Figure S8**), consistent with the inhibition mechanism being distinct from the impaired oligomer to fibril conversion observed for ΔP1 and ΔP2. Furthermore, we confirmed that these first 11 N-terminal residues are dispensable for oligomerization, as ΔN11 αS effectively assembles into WT-like oligomers under conditions used to generate kinetically trapped WT αS described above (**Figure S8**).

Overall, these data demonstrate an essential role of P1 and P2 in facilitating oligomer to fibril conversion and define the mechanistic foundation for PSMα3 inhibition of αS aggregation. Based on this knowledge, we suggest that molecules able to bind P1 and P2 in the oligomeric state could possess intrinsic anti-amyloidogenic properties by suppressing the oligomer to fibril transition.

### The familial G51D mutation impairs N-terminal mediated oligomer-to-fibril conversion and chaperone-assisted disaggregation

Familial mutations in the gene encoding αS often lead to a more aggressive form of Parkinson’s disease2. For instance, patients carrying the aS G51D mutation experience a more severe clinical course of the disease, characterized by earlier symptoms onset and significant psychiatric and autonomic dysfunctions^24,25^. The G51D mutation localizes to the P2 region, our findings suggesting that it may influence the conformational dynamics of the N-terminal domain and hence the ability of oligomers to convert into amyloid fibrils. This aligns with the previous observation that this amino acid change alters the oligomer conformation and induces a distinctive α-helical secondary structure component in their circular dichroism (CD) spectra^26^, which we corroborated here (**Figure S9**). To delve deeper into the impact of this mutation on the properties of the oligomers, we analyzed the differential deuterium uptake of WT and G51D aS oligomers. Compared with WT αS oligomers, G51D aS oligomers showed a significant increase in deuterium uptake in their N-terminal region involving residues 17-38 as exemplified by deprotection, impacting P1 (**Figure 4a** and **Figure S1**). This increased deuterium accessibility indicates a conformational transition N-terminal to the mutation site, implying a long-range effect in the oligomer structure and/or dynamics exerted by the G51D amino acid substitution.

**Figure 4.**
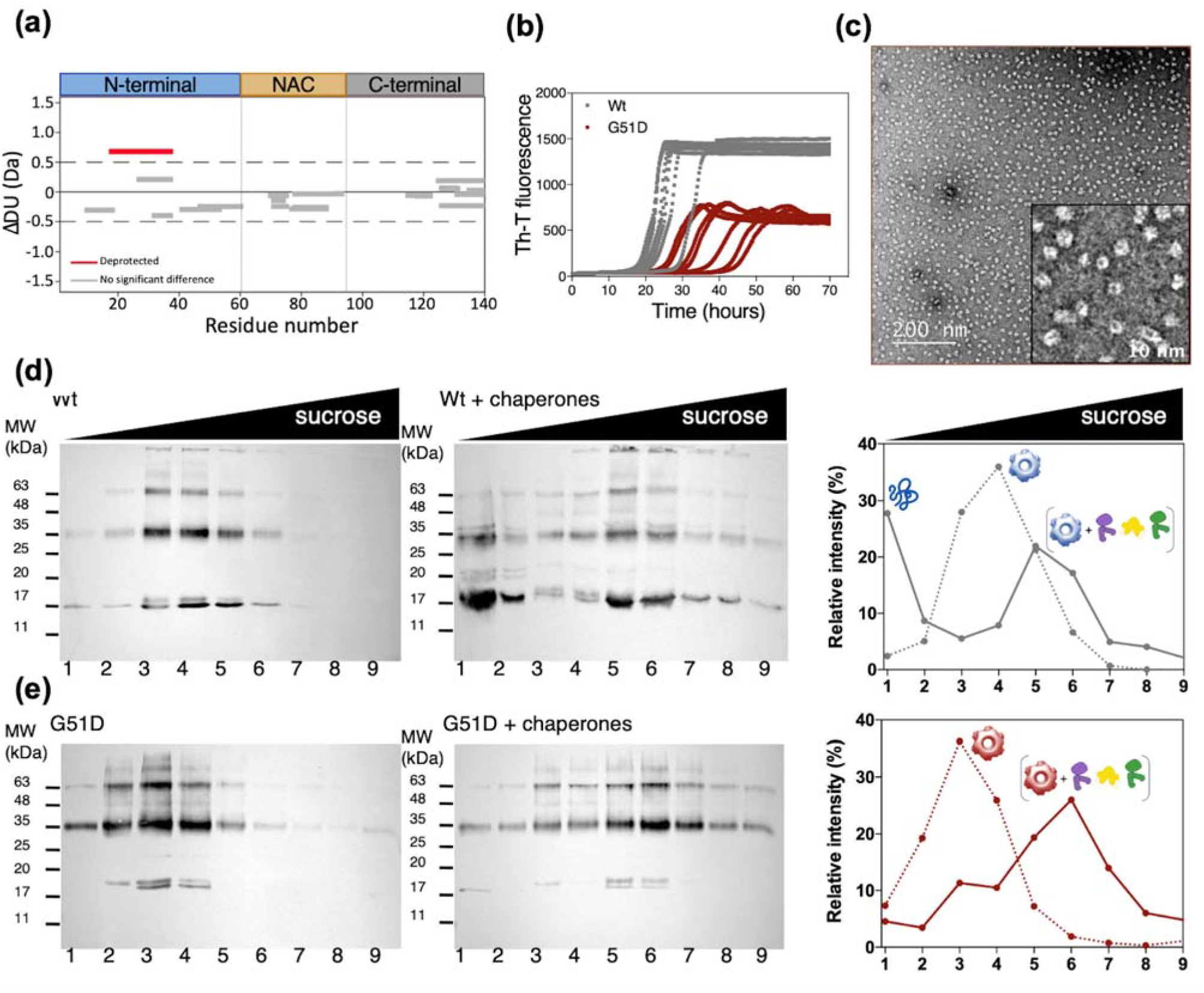
The effect of the familial G51D mutation on αS amyloid and oligomer formation and disaggregation by molecular chaperones. (a) Woods plot showing the relative solvent exposure/hydrogen bonding of G51D αS oligomers compared with WT αS oligomers by HDX-MS at the sixty second timepoint of exposure to deuterium. Deprotection from deuterium uptake occurs in the N-terminal region, as indicated by the peptide region colored in red. (b) Assembly kinetics of G51D into amyloid fibrils monitored using ThT fluorescence. (c) Representative nsEM micrographs of the G51D oligomeric fraction after 28 hours of assembly. (d and e) Sucrose-gradient fractionation of WT (d) and G51D (e) oligomers in the absence (left panels) or upon 2.5 hours incubation at 30 °C with the human disaggregase at αS:Hsc70 1:1.5 molar ratios (central panels). The distribution across the gradient was followed by western blot analysis using an anti-αS antibody. The intensity of the immunoreactive bands was quantified, and the graph (right panels) shows the relative intensities for non-treated (dotted line) and treated (solid line) oligomers.

The hereditary G51D mutation is also known to attenuate αS aggregation^27^, suggesting that the induced conformational rearrangement may also affect oligomer to fibril conversion. As observed for the synthetic mutants in P1 and P2, the delayed aggregation of G51D αS is associated with the accumulation of WT-like oligomers at the timepoint of maximal difference (t = 28 hours) compared with the WT fibril formation kinetics (**Figure 4b and c**). The G51D variant thus exemplifies how a disease-associated mutation in the P2 region elicits a structural rearrangement of the N-terminal region of the αS oligomer, including the P1 sequence, which further impacts oligomer to fibril conversion.

The N-terminal region affected by the G51D substitution identified above encompasses a well-established Hsc70 binding site (residues 37-43)^28,29^. Hsc70 is a fundamental element of the mammalian chaperone disaggregation machinery, working synergistically with its co-chaperones DNAJB1 and Apg2^29,30^. We decided to assess if the observed conformational differences in the Hsc70 binding site could alter its ability to be processed by chaperones. We hence monitored chaperone-mediated disaggregation of oligomers formed by WT and G51D αS using density-gradient centrifugation (**Figure 4d, e** and **Figure S10**). WT αS oligomers were efficiently disaggregated into monomers that floated to the top of the sucrose gradient at αS:Hsc70 1:1.5 molar ratios, while G51D oligomers were barely solubilized under the same conditions, exhibiting a greater resistance to Hsc70-mediated disaggregation (**Figure 4d and e**). Sample fractionation also revealed that the remaining oligomers of both proteins bound DNAJB1 and Hsc70, moving to more dense fractions of the sucrose gradient (**Figure 4d, e and Figure S10**). G51D oligomers are thus able to recruit the disaggregation machinery but they are not effectively processed; leading to an unproductive interaction where the G51D αS oligomers kidnap these essential protein quality control elements.

## DISCUSSION

The recent resolution revolution in cryo-EM has significantly advanced our structural understanding of the fibrillar amyloid state^31–33^. In contrast, the structure of intermediate oligomeric species remains largely uncharted^34^, hampering the development of oligomer-focused therapeutic strategies, despite their potential clinical relevance. For instance, Lecanemab, an FDA-approved monoclonal antibody, mitigates cognitive decline in Alzheimer’s disease by binding soluble Aβ aggregates (oligomers and protofibrils) with high selectivity over monomers^35^.

In this study, we characterized the binding site of PSMα3 within αS oligomers, leveraging this information to interrogate oligomer structural properties and the molecular determinants of oligomer progression to the amyloid state. We demonstrate that a specific motif involving residues 36-60 in the N-terminal of αS mediates PSMα3 selective binding to αS oligomers. This binding site has a distinct conformation in αS monomers and oligomers, a feature likely responsible for the oligomer-specific binding of PSMα3. Our data further reveal that this region populates a dynamic, yet defined, folded or partially folded conformation in the oligomer. Importantly, this N-terminal motif remains solvent accessible and targetable in the oligomeric state.

The PSMα3-mediated inhibition of oligomer to fibril conversion prompted us to investigate the role of its binding site, encompassing P1 and P2 regions, in this essential process of αS pathogenesis (**Figure 5**). Substantial deletions in both the N-terminal (ΔN11, ΔP1 and ΔP2 variants) and C-terminal (in references^22,36^) regions do not compromise oligomer formation, indicating that the NAC domain acts as the primary driver of oligomerization. Nevertheless, we found that the NAC region alone is insufficient to trigger the conversion of oligomers into fibrils, likely due to its rigidity and burial within the oligomer assembly. Instead, we show that the N-terminal regions that flank the αS NAC domain, involving particularly the P1 and P2 regions therein, modulate oligomer-to-fibril conversion in a sequence-dependent manner, facilitating the escape from the oligomer local thermodynamic minimal towards the fibrillar state.

**Figure 5.**
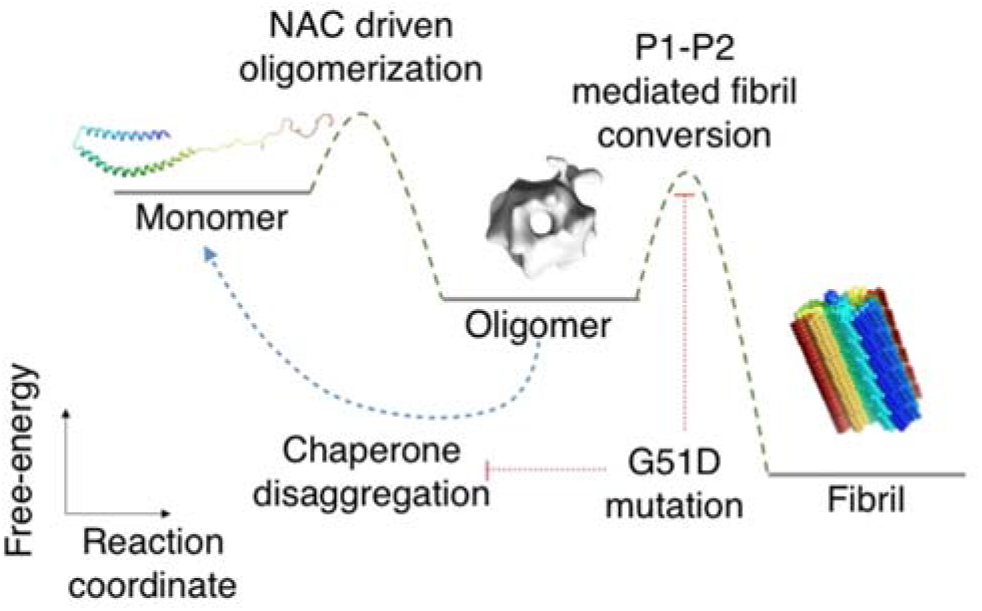
Schematic representation of the αS aggregation landscape.

Considering the oligomer-specific binding of PSMα3, along with its ability to inhibit oligomer to fibril conversion and mitigate neurotoxicity^16,^ targeting this specific region emerges as an appealing strategy for designing novel molecular entities mirroring PSMα3 activities. In previous endeavors^16^, we demonstrated that PSMα3 binding to oligomers is encoded in its physicochemical properties, not contingent on a specific sequence. In this way, sequentially divergent helical peptides with physicochemical traits akin to PSMα3, exhibited comparable inhibitory, binding, and neuroprotective properties^16^. This knowledge could potentially guide the development of a toolbox of sequentially diverse protein scaffolds mimicking PSMα3 properties to target αS oligomers^37^.

Our findings also provide a molecular framework to rationalize the impact of αS genetic mutations associated with familial Parkinson’s disease on oligomers. Most (but not all) reported familial mutations cluster at P2, having the potential to impact oligomer conformation, fibril conversion and interaction with other cellular components. Supporting this idea, the G51D amino acid substitution causes a change in oligomer structural dynamics in the N-terminal region, which delays the oligomer to fibril transition presumably results in an accumulation of long-lived, toxic^26^ oligomers that are not efficiently processed by the human disaggregase chaperone network and, instead, capture essential elements of this machinery. This evasion/impairment of proteostasis could explain why this αS mutation triggers the onset of Parkinson’s disease at ages when protein homeostasis is assumed to be preserved.

Finally, it is worth noting that PSMα3 binding reduces oligomer conformational heterogenicity, thereby improving the quality of the 2D image classification in cryo-EM, so we could unveil a previously unreported sixfold symmetry in the oligomer. Whereas limitations in resolution prevented a more detailed structural analysis, this is -to our knowledge-the first proof of a symmetrical supramolecular architecture in αS oligomers. Considering the low molecular weight of PSMα3 (2.6 kDa), it could be expected that larger ligands might exert a greater rigidification effect. Thus, our results encourage the use of P1 and P2 targeting molecules as a new route for advancing oligomer structural characterization.

## CONCLUSIONS

Our investigation identifies and characterizes a novel and pathologically relevant region for the implementation of oligomer-targeting strategies, while advancing our understanding of oligomer structural properties and the molecular mechanisms that underlie Parkinson’s disease pathogenesis.

## ASSOCIATED CONTENT

### Supporting Information

Materials and Methods, supplementary figures S1 to S12 and supplementary tables S1 to S4 (PDF).

### Data availability

The raw HDX-MS data have been deposited to the ProteomeXchange Consortium via the PRIDE/partner repository with the dataset identified PXD038573 (Reviewer account details: Username; Password: T14hHkoY). The mass spectrometry proteomics data have been deposited to the ProteomeXchange Consortium via the PRIDE/partner repository with the dataset identifier PXD039075 (Reviewer account details: Username: reviewer_; Password: 2lhLHYqS Y). The Cryo-EM maps for the 3DR a-syn C1, 3DR a-syn C6, 3DR a-syn:PSMa3 C1 and 3DR a-syn:PSMa3 C6 have been deposited at the Electron Microscopy Data Bank with accession codes EMBD-16466, EMBD-16527, EMBD-16525 and EMBD-16529 respectively.

## AUTHOR INFORMATION

### Author Contributions

The manuscript was written through contributions of all authors. / All authors have given approval to the final version of the manuscript. /

### Funding Sources

Spanish Ministry of Economy and Competitiveness (MINECO)

grant BIO2016-78310-R and BIO2017-91475-EXP (SV)

Spanish Ministry of Science and Innovation (MICINN) grant

PID2019-105017RB-I00 (SV)

MICINN grant PID2019-105872 GB-I00 (JMV)

MICINN grant PID2019-111068GB-I00 (AM)

ICREA-Academia 2015 (SV)

MICINN doctoral grant FPU17/01157 (JS)

Early postdoc mobility project SNSF P2EZP2_184258 (AL)

Basque Government grant IT1745-22 (FM)

BBSRC BB/M011151/1 grant (EJB)

Sir Henry Dale Fellowship jointly funded by Wellcome and the Royal Society 220628/Z/20/Z (ANC)

Royal Society grant RGS\R2\222357 (ANC)

University Academic fellowship from the University of Leeds (ANC)

Royal Society Research Professorship RSRP/R1/211057 (SER)

## ACKNOWLEDGMENT

Authors thank the cryoEM CNB-CSIC facility (CRIOMECORR project ESFRI-2019-01-CSIC-16), the services of the CNB-CSIC Mass Spectrometry facility, the Biomolecular mass spectrometry facility (UL) funded by the BBSRC (BB/M012573/1), the Microscopy Services (UAB), the Laboratori de Luminiscència i Espectroscòpia de Biomolècules (UAB) and the Biophysical and Structural Chemistry Platform at IECB, CNRS UAR 3033, INSERM US001. We thank James Ault for technical support with HDX-MS experiments and David Brockwell for illuminating discussions on the P1 region of αS.

