## Supporting information for "A targetable N-terminal motif orchestrates α-Synuclein oligomer to fibril conversion"

### MATERIALS AND METHODS

#### αS expression and purification

αS expression was performed using a pT7-7 plasmid encoding the αS gene in *Escherichia coli* BL21 (DE3) cells. Cells were grown in LB medium supplemented with 100 µg/mL ampicillin or in isotope-enriched M9 minimal media supplemented with $^{13}$C-glucose and $^{15}$N-ammonium chloride to obtain uniformly $^{13}$C$^{15}$N labelled samples. Protein expression was induced at an optical density of 0.6-0.8 (600 nm) with 1 mM isopropyl β-D-thiogalactopyranoside (IPTG) for 4 h. Cells were harvested by centrifugation and washed up by resuspension and centrifugation in PBS pH 7.4. The cell pellets were resuspended in 10 mL per culture liter in lysis buffer (50 mM Tris pH 8, 150 mM NaCl, 1 µg/mL pepstatin, 20 µg/mL aprotinin, 1 mM benzamidine, 1 mM PMSF, 1 mM EDTA, and 0.25 mg/mL lysozyme), lysed by sonication and centrifuged at 20,000 *g* for 30 min. The supernatant was boiled during 10 min at 95 °C and centrifuged again at 20,000 *g* for 30 min. The soluble fraction was treated with 136 µL/mL of 10% w/v streptomycin sulfate and incubated for 15 minutes. Upon centrifugation, soluble extracts were fractionated by adding 1:1 of saturated ammonium sulfate. The insoluble fraction was resuspended in 10 mL Tris 20 mM pH 8 per culture liter and dialyzed against Tris 20 mM pH 8. The dialyzed protein was filtered with a 0.22 µm filter and loaded onto an anion exchange column HiTrap Q HP (GE Healthcare, Chicago, USA). Tris 20 mM pH 8 and Tris 20 mM pH 8, NaCl 0.5 M were used as buffer A and buffer B. Fractions containing αS were further purified using size exclusion chromatography (Hiload 26/60 Superdex 75 preparation grade, GE Healthcare, Chicago, USA). Purified monomeric αS was dialyzed against 5 L ammonium acetate 50 mM in two steps; 4 h and overnight. Finally, protein purity was addressed using 15% SDS-PAGE. The purest fractions were lyophilized and stored at −80 °C. For the experiments, αS lyophilized aliquots were resuspended to a final concentration of 210 µM using PBS pH 7.4. αS concentration was determined by measuring the absorbance at 280 nm and using the extinction coefficient 5960 $M^{-1}$ $cm^{-1}$. For mutants lacking tyrosine 39, we used 4470 $M^{-1}$ $cm^{-1}$ as the extinction coefficient. All αS variants were purified under the same conditions.

#### Preparation of kinetically trapped αS oligomers and formation of the oligomer-PSMα3 complex

For the preparation of oligomeric samples, after the size exclusion step, purified αS was dialyzed against 5 L Milli-Q water and lyophilized for 48 h in aliquots of 6 mg. Aliquots were resuspended to a final concentration of 800 µM in PBS pH 7.4, filtered through 0.22 µm PVDF filters and incubated at 37 °C under quiescent conditions for 20–24 h. The incubated reaction was then ultracentrifuged at 288,000 *g* in a SW55Ti Beckman rotor, to remove fibrillar species formed during the incubation. The excess of monomeric protein was removed by four consecutive cycles of cleaning using 100 kDa centrifuge filters (Merck, Darmstadt, Germany). Oligomer concentration was measured using molar extinction coefficients determined using amino acid analysis (WT: 7000 $M^{-1}$ $cm^{-1}$; G51D: 12000 $M^{-1}$ $cm^{-1}$) in agreement with previous reports[1].

Oligomer-PSMα3 complex was prepared by incubating purified oligomer with a 3-fold molar excess of PSMα3 for 30 minutes. As previously reported, PSMα3 binds to αS oligomers with nanomolar affinity ($K_D$ = 6.67 nM) and with a 1:1 PSMα3: αS monomer ratio[2]. PSMα3 excess is removed by two consecutive cycles of cleaning with PBS pH 7.4 using 100 kDa centrifuge filters (Merck, Darmstadt, Germany).

**Solid-state NMR spectroscopy**

1.5 mg of $^{13}C$, $^{15}N$ labeled oligomer solution (PBS pH 7.4) was pipetted (approximately 50 µL) in a 4 mm Bruker rotor. All spectra were acquired in a 14.1 T Bruker magnet using a 4 mm Bruker HCN probe at 11 kHz MAS, 269 K probe temperature, resulting in sample temperature ~278 K. All 1D spectra were acquired with 64 scans with recycle delay of 3 sec. All cross-polarization based experiments were performed with a contact time of 600 us. The 2D 1H-13C INEPT spectra were acquired with 64 scans, 140 increments in indirect dimension, recycle delay of 2.2 sec, t1=7.7 ms, t2=18.4 ms resulting in a total acquisition time of 5.5 h. Both dimensions were apodized with Bruker QSine SSB=4 functions. The CP 2D 50 ms PDSD spectra was acquired with 512 scans, 300 increments in indirect dimension, recycle delay of 3 sec, t1=7.6 ms, t2=18.4 ms (with 90 kHz $^{1}H$ SPINAL-64 decoupling) resulting in total acquisition time of 128 h. Both dimensions were apodized with Bruker QSine SSB=3 functions. Water-edited experiments were recorded using a $T_2$ filter of 1.5 ms and a variable $^{1}H$-$^{1}H$ spin diffusion period, using 1024 scans. Results were plotted as the square root of the $^{1}H$-$^{1}H$ mixing period. The spectra were processed and analyzed using TopSpin3.61 and CcpNmr version 2.4.1 programs.

**Negative staining electron microscopy**

For negative staining electron microscopy analysis, samples were diluted to a concentration of 0.2-0.05 mg/mL in PBS and placed onto glow-discharged carbon-coated copper grids to adsorb for 1 min. The excess of sample was carefully blotted using ashless filter paper. Grids were negatively stained with 2% (w/v) uranyl acetate for 1 min and excess of uranyl acetate was absorbed using ashless filter paper. A TEM JEOL JEM1400 microscope was used operating at an accelerating voltage of 120 kV equipped with a CCD GATAN 794 MSC 600HP camera.

Representative images of each grid were selected. For particle analysis and 2D classification, images were acquired using a JEOL JEM 1010 electron microscope operated at 100 kV and equipped with a CCD camera (4 K × 4 K TemCam-F416, TVIPS). Images were recorded at a 50,000 × nominal magnification with a sampling rate of 2.4 Å/px. These images were processed following the Scipion3 processing workflow[3]. Images were CTF-corrected using CTFFIND4[4]. Particles were automatically selected using Xmipp3[5] and 2D-classified using Relion2[6] and CryoSPARC[7].

**CryoEM data acquisition**

Aliquots of 4 µL of αS oligomers and αS oligomers-PSMα3 complexes were vitrified using a Vitrobot Mark IV (FEI) and were incubated onto Quantifoil R 2/2 300 mesh grids with an additional ultrathin continuous carbon layer, blotted for 2 s at 22 °C and 95% humidity and plunged into liquid ethane. The cryoEM grids were checked and data from the best one was acquired in a 200 kV FEI Talos Arctica equipped with a Falcon III direct electron detector at the Centro Nacional de Biotecnología (CNB) cryoEM facility. A total of 967 movies for αS oligomers and 852 movies for αS oligomers-PSMα3 complexes were acquired at a nominal magnification of 73,000x (corresponding to a pixel size of 1.37 A/pix), with a defocus range of 1.4 to 3.2 µm. Exposure was set to 0.9322 e-/Å/sec and 30 frames were collected in total, with an overall dose of 28 e-/Å for αS oligomers and 1.06 e-/Å/sec and a total dose of 32 e-/Å for αS oligomers-PSMα3 complexes.

**Image processing and three-dimensional reconstruction**

Image processing of αS oligomers and αS oligomers-PSMα3 complexes was performed following a similar workflow. All programs used for image processing to obtain the different 3D maps are implemented in the Scipion software platform. First, the movies were aligned using MotionCor2[8] and the outputs were subjected to CTF determination using Gctf[9]. Particles were automatically picked with Xmipp3 –auto-picking software. The 193,427 (αS oligomers) and 187,446 (αS oligomers-PSMα3 complex) extracted particles were subjected to several 2D classifications using Relion 2.0 and Cryosparc to exclude bad particles and ice contamination. Some of the best 2D classes were used as a template to generate an initial model using both Cryosparc and RANSAC[10]. In both cases, models were low-pass filtered to 50 Å and used for a 3D classification of 85,628 (αS oligomers) and 76,730 (αS oligomers-PSMα3 complex) particles contained in the best 2D classes performed without symmetry imposition. The particles of the best classes were used for a further 3D auto-refine using Relion 2.0 and yielded cylindrical oligomeric structures at 18.5 Å (αS oligomers) and 19 Å (αS oligomers-PSMα3 complex) resolution (**Supplementary Table 3**).

**Small-angle x-ray scattering data acquisition**

SAXS measurements were performed on an in-house instrument at Aarhus University. The instrument is a modified NanoSTAR from Bruker AXS with a homebuilt scatterless pinhole in front of the sample and an Excillum liquid metal jet source[11]. αS oligomers at a concentration of 4.6, 2.0, and 1.3 mg/mL were measured in the same flow-through capillary. Scattering from the buffer was measured and subtracted as background and the data were converted to absolute intensity scale using the scattering from a pure water sample as standard[12]. The azimuthally averaged intensity $I(q)$ was calculated from the two-dimensional data as a function of the modulus of the scattering vector $q$. The data were displayed in Guinier plots of $\ln(I(q))$ vs $q^2$ and Kratky plots of $q^2\, I(q)$ vs $q$ to check for, respectively, aggregation and flexibility. The former also gives the radius of gyration $R_g$ and the forward scattering $I(0)$. An Indirect Fourier Transformation (IFT) of the data[13] was used for obtaining the pair distance distribution function, $p(r)$, which is a histogram of distances between pairs of points within the structure.

**Modelling of Small-angle x-ray scattering data**

A model based on super-ellipsoid of revolution was constructed[14]. It consists of a super-ellipsoid core with a cylindrical hole along the symmetry axis of super-ellipsoid surrounded by a shell with a different (lower) density and constant width. Both the outer surface and the core-shell interface were graded by including Gaussian factors as done by Maric et al.[14]. This publication also contains the equations required for calculating the present model scattering. The shape parameter of the super-ellipsoid was fixed at t = 4 for both core and shell. The scattering from the internal structure of the shell with the random coils were included as described in ref[15] using an extra term with the form factor of the shell subtracted from the random coil scattering. The latter was described by the scattering from Gaussian chains multiplied by a cross-section Guinier term $\exp(-R_c^2 q^2/4)$, where $R_c$ is the cross-section radius[16]. The intensity of the models is expressed on absolute scale using a excess scattering density of $2.00 \times 10^{-10}$ cm/g for the protein. The monomer was set to 14.5 kDa for αS.

The model depends on the following parameters. The core radius $R$, the axis ratio $\varepsilon$, the grading width of the core-shell interface $\sigma_{in}$, the radius of the hole in the core $R_{hole}$, the width of the shell $W_{shell}$, the grading width of the outer surface of the shell $\sigma_{shell}$, where the restraint $W_{shell} = 2\,\sigma_{shell}$ was used. Additional parameters are relative density of the shell $\rho_{shell}$, the radius of the random coils chains $R_c$ and the aggregation number $N_{agg}$. The fraction of protein in core $f_{core}$ and shell (1 - $f_{core}$) can be calculated from the other fit parameters. The mass of a protein chain in the shell was used for calculating the radius of gyration $R_g$ for the random coil chains scattering using an expression for unfolded polypeptide chains[17].

In a log-log plot, the data sets (**Supplementary Figure 11A**) are nearly identical for the different concentrations except for scale factors. The data display a crossover towards constant intensity as $q$ goes to zero. At higher q, it is followed by a power-law behaviour with an exponent of approximately –4. At even higher $q$, there is a shoulder with a subsequent power law with a lower exponent, in agreement with the presence of some polymer-like scattering due to the presence of some random coils in the structure.

Guinier plots of the data a ln($I$($q$)) vs $q^2$ at the highest concentrations (**Supplementary Figure 11B**) show a linear behaviour. The radius of gyration is $R_g$ = 76.1 ± 0.3 Å and the forward scattering $I$(0) = 1.31 ± 0.01 $cm^{-1}$. An Indirect Fourier Transformation (IFT) of the data[3, 4] gave the functions and the fits displayed in (**Supplementary Figure 11C**) for a maximum distance of 265 Å. The IFT gave as values $R_g$ = 80.5 ± 0.2 Å and $I$(0) = 1.384 ± 0.003 $cm^{-1}$, deviating slightly from the values determined by the Guinier plot and fit. Kratky plots of $q^2$ $I$($q$) vs $q$ (**Supplementary Figure 11D**) have a maximum at low $q$ in agreement with a compact globular-like structure and levels off at large $q$ in agreement with additional presence of random coil scattering.

To display the resulting structures, a program was written that generates dummy atoms with a density that follows that determined by the modelling[14]. The resulting dummy atoms and coordinates are saved in PDB format. The density of the dummy atoms represents the ensemble average, so the chain structures are not present in the dummy atom representations.

**Crosslinking experiments and mass spectrometry analysis (XL-MS)**

30 µg of αS oligomers and αS oligomers-PSMα3 complexes were subjected to chemical crosslinking by incubation with 15 mM DMTMM in PBS pH 7.4 for 30 min at RT. The reactions were quenched for 15 min at RT by adding 50 mM Tris-HCl pH 7.0. DMTMM-crosslinked samples were incubated in Laemmli sample buffer (0.02% [w/v] bromophenol blue, 2% [w/v] SDS, 10% [v/v] glycerol, 60 mM Tris-HCl pH 6.8) for 5 min at 96 °C and loaded onto a 12% polyacrylamide gel. The gel was stained, and the visible bands were excised and subjected to automated reduction, alkylation with iodoacetamide and trypsin digestion in a Proteineer DP robot (Bruker Daltonics). The resulting peptide mixture was speed-vac dried and re-dissolved in 0.1% (v/v) formic acid. Liquid chromatography–mass spectrometry (LC-MS/MS) analysis was carried out

using a nano-LC Ultra HPLC (Eksigent, Framingham, MA) coupled online to a 5600 triple TOF mass spectrometer (AB Sciex, Framingham, MA) through a nanospray III ion source (AB Sciex) equipped with a fused silica PicoTip emitter (10 µm x 12 cm; New Objective, Woburn, MA). Peptides were fractionated at a flow rate of 0.250 mL/min at 50 °C under gradient elution conditions. The ion source was operated in positive ionization mode at 150 °C with a potential difference of 2300 V.

For peptide identification, raw MS data was searched against a custom-made database containing the amino-acid sequence of human αS or PSMα3. The MS/MS ion search was performed with MeroX 2.0[18]. Search parameters were set as follows: DMTMM as crosslinker and trypsin as enzyme, allowing 3 missed cleavages for Arg and Lys. Carbamidomethylation (Cys) and oxidation (Met) were set as fixed and variable modifications, respectively. Analysis was performed with MS and MS/MS tolerances of 10 and 20 ppm, respectively. Peptide identifications were filtered at an FDR < 5% and an XlinkX score > 30 and all the MS2 spectra of the resulting peptides were manually revised.

**Hydrogen-deuterium exchange-mass spectrometry (HDX-MS)**

HDX-MS experiments were performed using an automated HDX liquid handling robot (LEAP Technologies, Ft Lauderdale, FL, USA) coupled to an Acquity M-Class LC and HDX manager (Waters, UK). Samples contained 50 µM of αS monomer or oligomer in PBS buffer, pH 7.4. The robot was used to transfer 95 µL of deuterated buffer (PBS, pD 7.4, 0.01% w/v DDM) to 5 µL of protein-containing solution and the mixture was incubated at 4°C for 0, 0.5, 1 or 5 min. Three replicate measurements were performed for each time point and for each protein condition. 75 µL of quench buffer (PBS buffer, 4 M guanidine HCl, 0.05% w/v DDM, pH 2.1) was added to 75 µL of the labelling reaction to quench deuterium exchange. 50 µL of the quenched sample was injected into an Enzymate immobilised pepsin column (Waters, UK). A VanGuard Pre-column [Acquity UPLC BEH C18 (1.7 µm, 2.1 mm x 5 mm, Waters, UK)] was used to trap the peptides produced for 3 min. A C18 column (75 µm, 2.1 mm x mm, Waters, UK) separated peptides using a gradient of 0-40% (v/v) acetonitrile (0.1% v/v formic acid) in $H_2O$ (0.3% v/v formic acid) over 7 min at 40 µL $min^{-1}$. Separated peptides from the LC column were infused into a Synapt G2Si mass spectrometer (Waters, UK) operated in $HDMS^E$ mode. Peptides were separated by ion mobility prior to CID fragmentation in the transfer cell for peptide identification. Deuterium uptake was quantified at the peptide level.

Data analysis was performed using PLGS (v3.0.2) and DynamX (v3.0.0) (Waters, UK). Search parameters in PLGS were: peptide and fragment tolerances: automatic, minimum fragment ion matches: 1, digest reagent: non-specific, false discovery rate: 4. Restrictions for peptides in DynamX were: minimum intensity: 1000, minimum products per amino acid: 0.3, maximum sequence length: 25, maximum error = 5 ppm, file threshold: 3. Peptides with significant increase/decrease in deuterium uptake were identified using a hybrid significance test implemented in Deuteros (Table S4)[19]. Sequence coverage is summarized in **Figure S12**. Woods plots were generated using Deuteros[19].

**Assembly kinetics into amyloid**

αS amyloid aggregation was monitored in non-binding 96 well plates (Corning, Corning, NY, USA). Each 100 µL reaction contained 100 µM αS and 20 µM thioflavin-T (Th-T) in PBS pH 7.4. Aggregation kinetics were recorded using a Spark plate reader (Tecan, Switzerland) at 37 °C and 600 rpm with a time interval of 15 minutes (Ex. 445 nm, Em: 495 nm).

**Isolation of low molecular weight aggregates generated during αS in vitro aggregation**

αS aggregation was performed as described in the previous section. For *wild-type*, ΔN11, ΔP1, ΔP2, ΔΔ and Y39A endpoint samples were recovered for the plate, whereas for S42A and G51D variants, we recovered the samples at the time point of maximal inhibition (28 hours). To isolate the oligomeric fraction, we adapted the centrifugation-based protocol developed by Kumar and coworkers[20]. αS preparations (400 µL) were subjected to ultracentrifugation at 100,000 g for 60 min at 20 °C in a SW55Ti Beckman rotor to remove larger fibrillar species. The soluble fraction was then filtrated through 100 kDa centrifuge filters (Merck, Darmstadt, Germany) to fractionate low molecular weight aggregates and monomeric αS. The filtrated sample contains monomeric αS, whereas oligomers are retained in the upper section of the filter in a volume of 30-50 µL. The monomer excess was then washed by diluting the sample in PBS pH 7.4 to 500 µL and the procedure repeated twice. The oligomeric fraction is then recovered by carefully pipetting (ca. 30-50 µL) and was subsequently analyzed by transmission electron microscopy as previously described above. This procedure allows a morphological characterization of low molecular weight aggregates generated in the aggregation reaction by concentrating these low populated species and removing the large excess of monomer that would preclude their visualization by EM in an unprocessed sample.

**Far circular dichroism analysis**

Far-UV CD spectra of the oligomer preparations were recorded on a Jasco J-815 CD spectrometer (Halifax, Canada) at 25 °C. Oligomer concentration was adjusted to 5 μM in PBS pH 7.4. CD signal was measured from 260 nm to 190 nm at 1 nm bandwidth, 1 sec of response time and a scan speed of 200 nm/min on a 0.1 cm quartz cell. Ten to twenty accumulations were recorded and averaged for each measurement.

**Chaperone disaggregation**

Hsc70, DNAJB1 and Apg2 chaperones were produced as previously reported[21]. Products of the disaggregation reaction were fractionated in a sucrose gradient as follows. Samples (400 μL) containing 10 μM αS oligomers, 10 μM Hsc70, 5 μM DNAJB1 and 1 μM Apg2 were incubated 2.5 hours in 40 mM Hepes-KOH pH 7.6, 50 mM KCl, 5 mM MgCl2, and 2 mM DTT at 30 °C, buffer that also contained ATP (2 mM) and the ATP-regeneration system (8 mM phosphoenol pyruvate and 20 ng/μL pyruvate kinase). Reaction mixtures were then applied to 3.2 mL of a 5–40 % sucrose gradient and centrifuged at 162,000 g for 2 hours at 4 °C. Fractions (400 μL) were manually removed and subjected to SDS-PAGE followed by immunoblotting with anti-αS antibody (Invitrogen PA5-85343, 1:2000 dilution).

**SUPPLEMENTARY FIGURES**

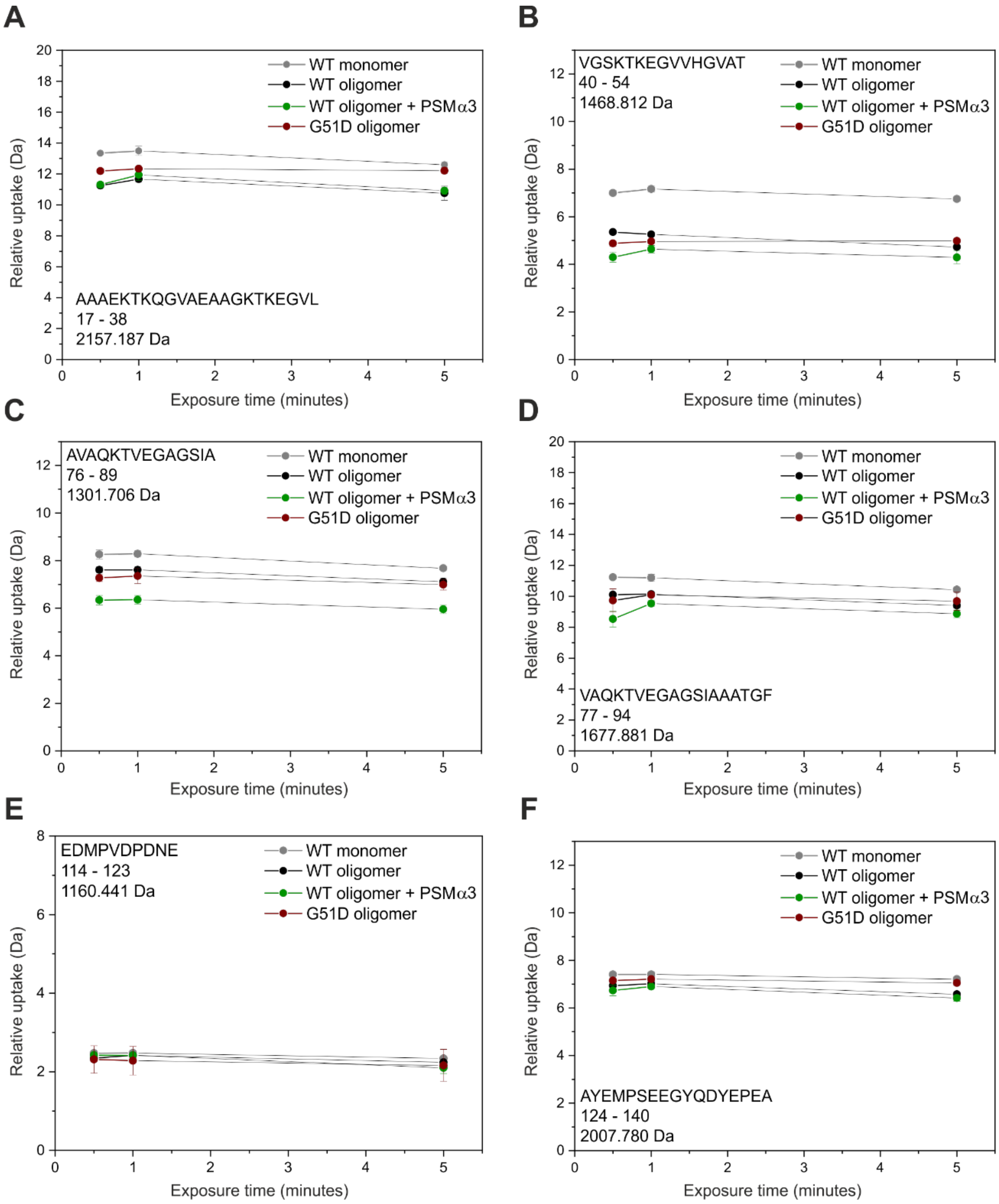


**Figure S1. The N-terminal and NAC regions of αS display significant protection from deuterium uptake in oligomeric states by HDX-MS**. Example deuterium uptake plots for residues (**A**) 17-38, (**B**) 40-45, (**C**) 76-89 (**D**) 77-94, (**E**) 114-123 and (**F**) 124-140 across 30 s, 1 min and 5 min exposure timepoints. WT monomer is shown in grey, WT oligomer in black, oligomers in the presence of PSMα3 in green and G51D oligomers in red. Protection from deuterium incorporation occurs across residues in the N-terminal and NAC regions in the WT oligomeric states compared to monomer. Binding of PSM3 increases the degree of protection observed. The degree of protection observed in the G51D oligomer is comparable to WT oligomers (in the absence of PSMα3). Each uptake plot is annotated with the peptide sequence, residue numbers and peptide mass in Da.

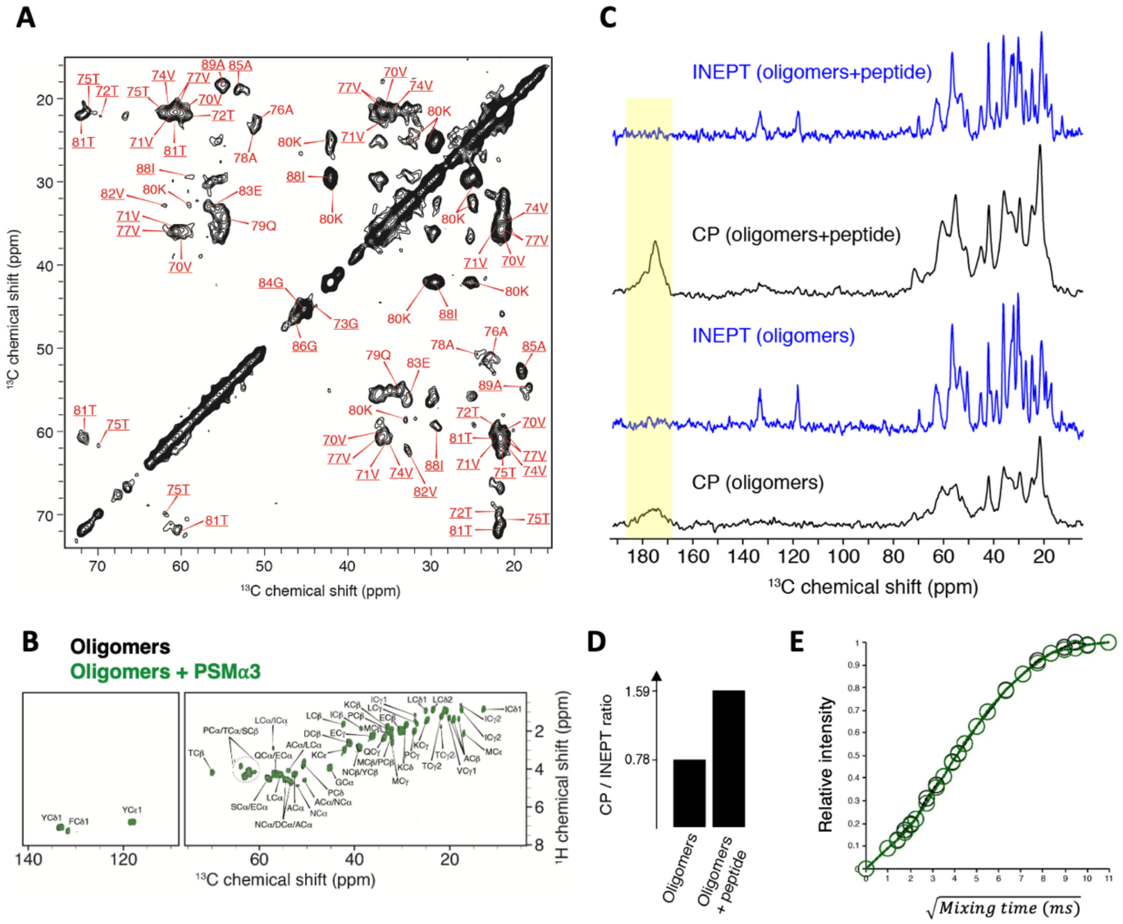


**Figure S2. MAS solid-state NMR analysis of $^{13}$C-labelled αS oligomers and oligomers in complex with PSMα3.** (**A**) 2D $^{13}$C-$^{13}$C PDSD correlation spectrum (mixing time of 50 ms) of oligomers. Residues were assigned based on a previous study of αS fibrils (PDB: 2N0A)[1]. Residues underlined have a chemical shift difference between oligomers and fibrils as $\Delta C\alpha < 1$ppm and $\Delta C\beta < 2$ppm. Residues not underlined as $1 < \Delta C\alpha < 2$ppm and $\Delta C\beta < 2$ppm. Spectra correspond to 1.5 mg of $^{13}$C, $^{15}$N labeled oligomer solution in PBS pH 7.4 in the absence or presence of PSMα3. Oligomer-PSMα3 complex was prepared as described in the Materials and Methods section. (**B**) 2D $^{1}$H-$^{13}$C INEPT spectra of oligomers (black) and oligomers + PSMα3 peptide (green). (**C**) 1D $^{13}$C spectra of oligomers and oligomers + PSMα3 peptide, detected using cross-polarization (CP) and INEPT as an initial $^{1}$H to $^{13}$C polarization transfer. The carbonyl spectral region is highlighted in pale yellow. (**D**) Ratio of CP/INEPT peak intensities integrated for the spectral region 0-75 ppm. (**E**) Water-edited analysis of the oligomers (black) and oligomers + PSMα3 peptide (green) samples.

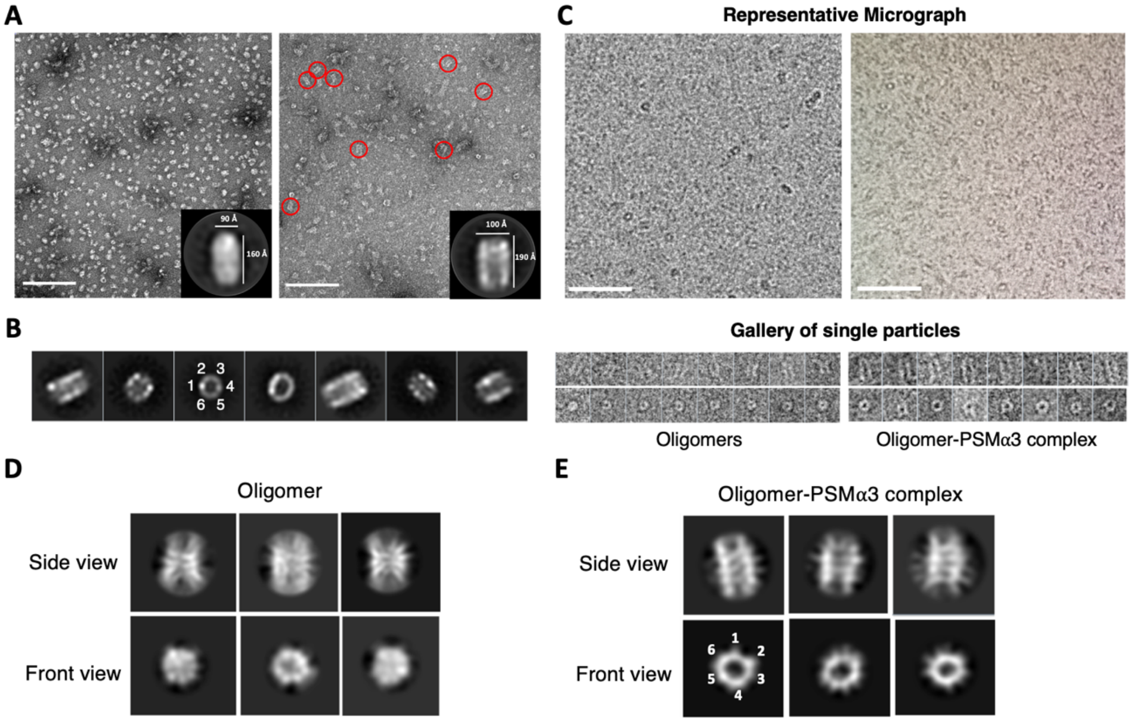


**Figure S3**. **Electron microscopy characterization of αS oligomers in the absence or presence of PSMα3.** (**A**) Representative EM micrographs from negatively-stained αS oligomers (left) and oligomer:PSMα3 complexes (right). Insets show a representative 2D class to illustrate PSMα3 induced rigidification. Scale bar: 100 nm. (**B**) Representative 2D classification of particles collected in panel **A**. (**C**) Representative micrographs and gallery of single particles obtained from the cryoEM analysis of αS oligomers in the absence or presence of PSMα3. Scale bar: 100 nm. (**D-E**) 2D cryoEM classes of oligomers (**D**) and oligomers in complex with PSMα3 (E). The observed symmetry is labeled with numbers.

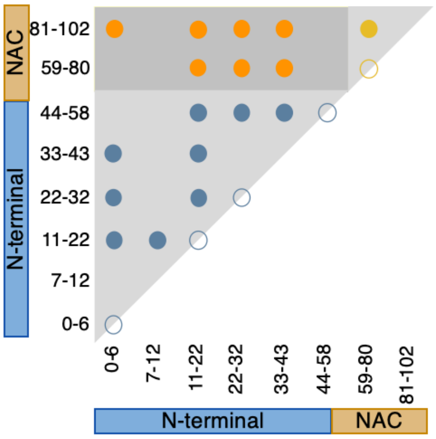


**Figure S4. Mapping structural contacts in αS oligomers by XL-MS**. Illustration of the N-terminal domain (blue), NAC domain (gold) and N-terminal to NAC (orange) contacts identified by crosslinking αS oligomers with DMTMM. Open circles represent crosslinks between identical fragments (necessarily intermolecular).

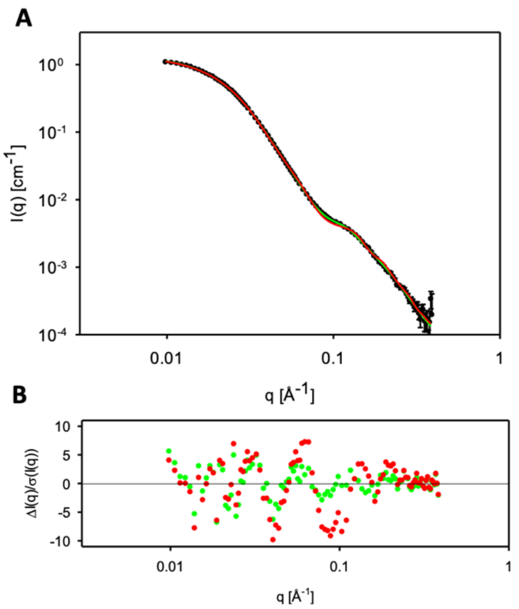


**Figure S5. SAXS analysis of oligomer organization**. (**A**) IFT fits to the SAXS data. The data for the αS oligomers are shown as black points. The fit for a model with one random coil chain per monomer is shown as the green curve and the fit for a model with two random coil chains per monomer is shown as a red curve. The two models have reduced chi-square of, respectively, $\chi^2$ = 5.9 and $\chi^2$ = 17.3, demonstrating a better fit with only one chain per monomer. (**B**) Residual of the difference between model and experimental intensities divided by the standard error on the data from counting statistics, colored as in (a).

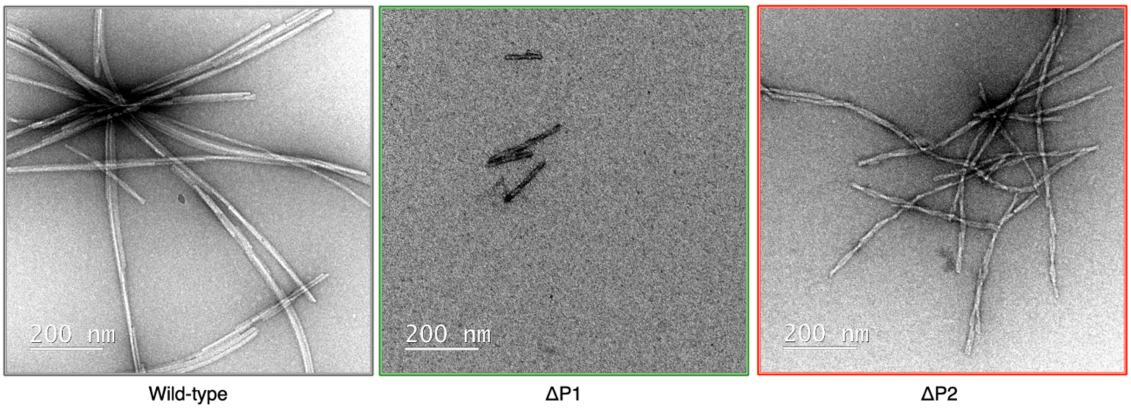


**Figure S6. Representative nsEM micrographs of WT, ΔP1 and ΔP2 variants at the end point of the aggregation reaction.** For ΔP1 and ΔP2 variants most of the grid area was devoid of fibrils. We show representative images of those observed.

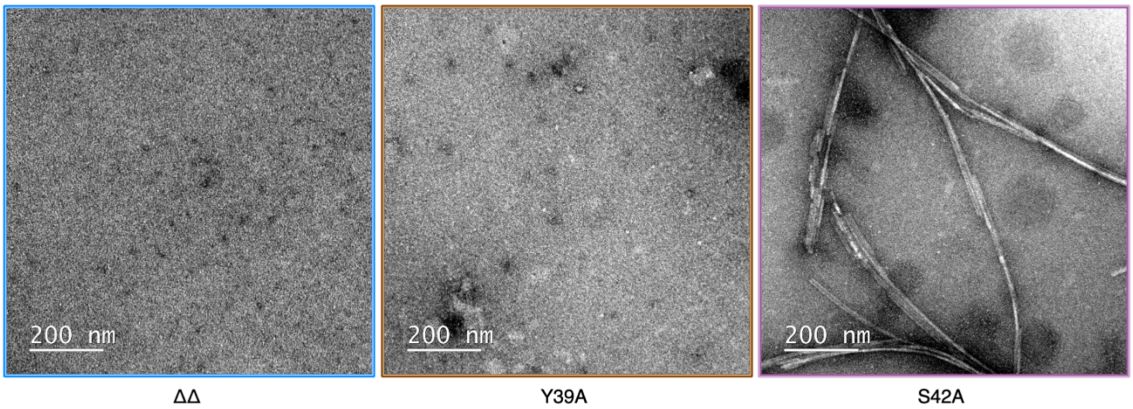


**Figure S7. Representative nsEM micrographs of ΔΔ (left), Y39A (center) and S42A (right) variants at the end point of the aggregation reaction.**

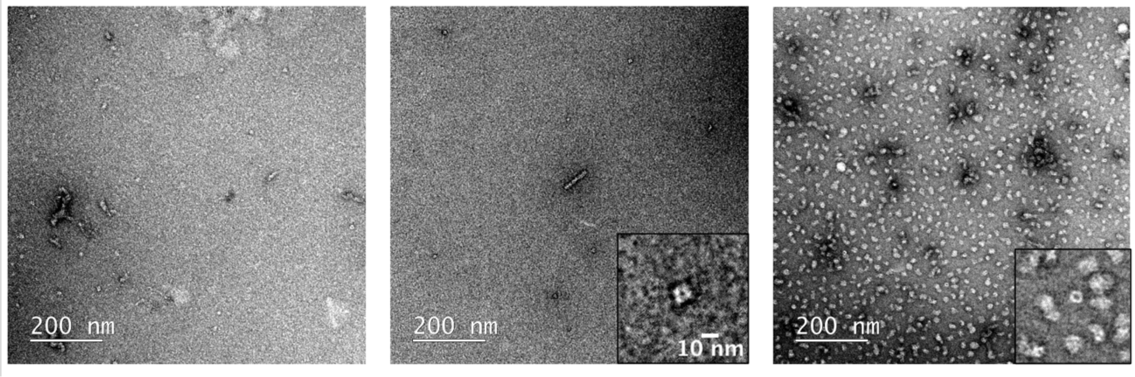


**Figure S8. Analysis of amyloid and oligomer formation of the ΔN11 αS variant.** Left: Representative nsEM micrograph at the endpoint of the aggregation reaction. Center: Representative nsEM micrograph of the oligomeric fraction isolated at the endpoint of assembly. Right: Representative nsEM micrographs of ΔN11 oligomers generated under oligomer forming conditions (kinetically trapped).

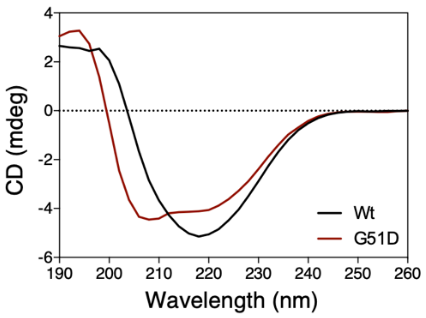


**Figure S9. Circular dichroism spectra of kinetically trapped oligomers of G51D (dark red) and WT αS (black).**

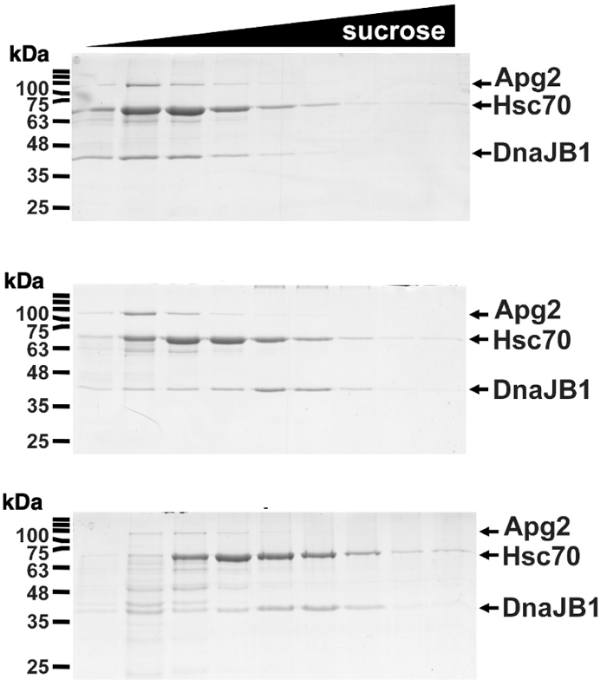


**Figure S10. Migration of the human disaggregase components in the sucrose gradient.** Distribution of the different chaperones in the absence of oligomers (upper panel) and in the presence of WT (intermediate panel) or G51D (bottom panel) oligomers was analyzed by SDS-PAGE and Coomassie staining. Samples correspond to those in Figure 3d and 3e, where 10 µM of Wt oligomers or G51D oligomers were incubated with 10 µM Hsc70, 5 µM DnaJB1, and 1 µM Apg2 at 30 °C in the presence of ATP and an ATP-regeneration system.

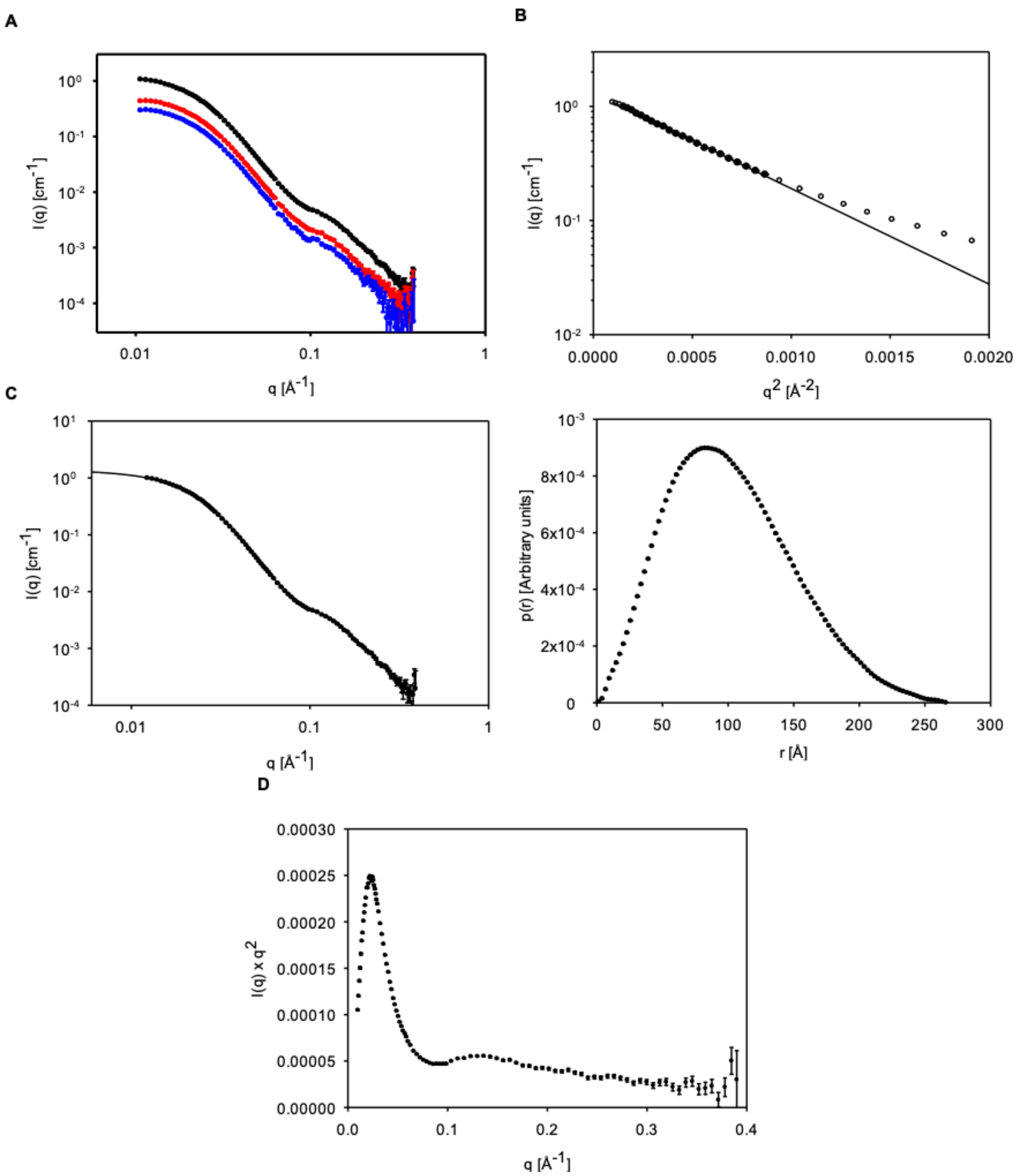


**Figure S11. SAXS measurements and data analysis.** (**A**) Concentration series for αS oligomer. From top to button: 4.6, 2.0, 1.3 mg/mL. (**B**) Guinier plots of the data for the high concentration samples of αS. Only the data points filled with black have been used for the fit (straight line). (**C**) Left: IFT fits to the data (lines) for the high concentration samples of αS (points). Right: Pair distance distribution functions for αS. (**D**) Kratky plots of the data for the high concentration samples of αS.

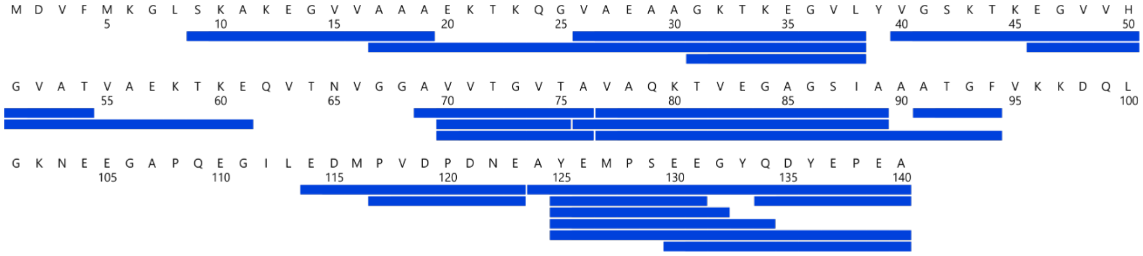


**Figure S12. Sequence coverage of αS monomers, oligomers, oligomers with PSMα3 and G51D oligomers used to detect deuterium uptake by HDX-MS.**

### SUPPLEMENTARY TABLES

**Table S1. $^{13}$C chemical shift differences between αS fibrils (BMRB entry 25518)[1] and oligomers.**

| Residue | ΔCα (ppm) | ΔCβ (ppm) |
|---|---|---|
| 70V | <0.1 | <0.1 |
| 71V | <0.1 | <0.1 |
| 72T | <0.1 | <0.1 |
| 73G | -0.3 | |
| 74V | <0.1 | <0.1 |
| 75T | <0.1 | 0.3 |
| 76A | -1.2 | -0.9 |
| 77V | <0.1 | <0.1 |
| 78A | -1.3 | 0.9 |
| 79Q | -2.4 | -0.7 |
| 80K | 1.2 | -0.3 |
| 81T | <0.1 | 0.5 |
| 82V | -0.5 | 1.2 |
| 83E | -2.2 | -0.4 |
| 84G | <0.1 | |
| 85A | 0.5 | -0.7 |
| 86G | <0.1 | |
| 87S | not detected | not detected |
| 88I | 0.7 | -1.7 |
| 89A | <0.1 | 0.5 |

**Table S2. Parameter values obtained from the SAXS analysis. The parameters are described in the methods section.**

| Parameter | αS oligomers |
|---|---|
| $N_{agg}$ | 31.5 ± 0.1 |
| $R$ [Å] | 42.7 ± 1.7 |
| e | 1.58 ± 0.05 |
| $s_{core}$ [Å] | 9.6 ± 1.8 |
| $W_{shell}$ [Å]calculated | 52.2 |
| $R_{hole}$ [Å] | 5.7 ± 1.7 |
| $s_{shell}$ [Å] | 26.6 ± 0.4 |
| $r_{shell}$ | 0.114 ± 0.008 |
| $f_{core}$ calculated | 0.519 |
| $R_c$ [Å] | 7.9 ± 0.5 |
| $R_g$ [Å] calculated | 15.9 |

**Table S3. CryoEM data acquisition and processing**

| Data collection & processing | αS oligomers | αS oligomers-PSMα3 |
|---|---|---|
| Microscope | FEI Talos Arctica | FEI Talos Arctica |
| Voltage (KV) | 200 | 200 |
| Camera | Falcon 3 | Falcon 3 |
| Total dose ($e^-/Å^2$) | 28 | 32 |
| Frames | 30 | 30 |
| Dose per frame ($e^-/Å^2$) | 0.932 | 1.06 |
| Defocus range | -1.4 to -3.2 | -1.4 to -3.2 |
| Pixel size (Å) | 1.37 | 1.37 |
| Initial number of particles | 193,427 | 187,446 |
| Final number of particles | 38,069 | 26,658 |
| FSC threshold | 0.143 | 0.143 |
| Symmetry | C1 | C1 |
| Map resolution (Å) | 18.5 | 19 |

**Table S4. HDX data summary table. SD = standard deviation, CI = confidence interval.**

| Data collection & processing | WT monomer | WT oligomer | WT oligomer +PSMα3 | G51D oligomer |
|---|---|---|---|---|
| **HDX reaction details** | PBS in 100% D2O, pH 7.4, 4°C | | | |
| **HDX time points (mins)** | 0, 0.5, 1, 5 | | | |
| **HDX controls** | Fully deuterated controls were not performed | | | |
| **Back-exchange** | N/A | | | |
| **No. of peptides** | 22 | | | |
| **Sequence coverage** | 75.0% | | | |
| **Average peptide length / redundancy** | 11.27 / 2.36 | | | |
| **Replicates** | 3 (Technical) | | | |
| **Repeatability** | 0.0735 (average SD) | | | |
| **Significant difference in HDX (ΔHDX > XD)** | 98% CI: 0.50 Da / p-value < 0.02 | | | |
